# Engineering a PD-L1–sensing synthetic receptor for programmable macrophage-mediated phagocytosis

**DOI:** 10.64898/2026.01.21.700810

**Authors:** De Martino Ilaria, Russo Luigi, Marchetti Matteo, Siciliano Velia

## Abstract

Macrophages are abundant immune cells within the tumor microenvironment with intrinsic phagocytic capabilities, yet their antitumor functions are frequently suppressed by inhibitory signals. Synthetic biology enables the rational design of ligand-responsive genetic circuits to reprogram immune cell behavior. Here, we report the engineering of a synthetic Notch–based receptor that detects PD-L1, an immune-checkpoint broadly expressed by cancer cells. Upon PD-L1 engagement, the circuit triggers programmable outputs, including expression of a fluorescent reporter or CV1-Fc, that locally blocks the CD47 “don’t eat me” signal. We show that circuit activation scales with PD-L1 levels, partially attenuates PD-1/PD-L1 signaling, and that conditional CV1-Fc expression enhances engulfment of SKOV-3 ovarian cancer cells by THP-1–derived macrophages in vitro. Collectively, this work reframes PD-L1 from an end-point therapeutic target to a primary input signal for synthetic circuit activation and establishes a modular framework for engineering macrophage behaviour through spatially confined, ligand-responsive control.

**Graphical abstract. A novel α-PDL1 SNIPR receptor to program macrophage outputs.** PD-L1 expressed on cancer cells is detected by α-PD-L1 SNIPR–engineered macrophages, triggering release of the GAL4-VP64 transcription factor and activation of a programmable actuator. Circuit activation leads to customized outputs, including biosensing and enhanced macrophage phagocytosis via CD47 blockade, as well as immunomodulatory effects through interference with the PD-1/PD-L1 checkpoint axis. This sensor–actuator framework enables spatially confined and ligand-dependent reprogramming of macrophage function.

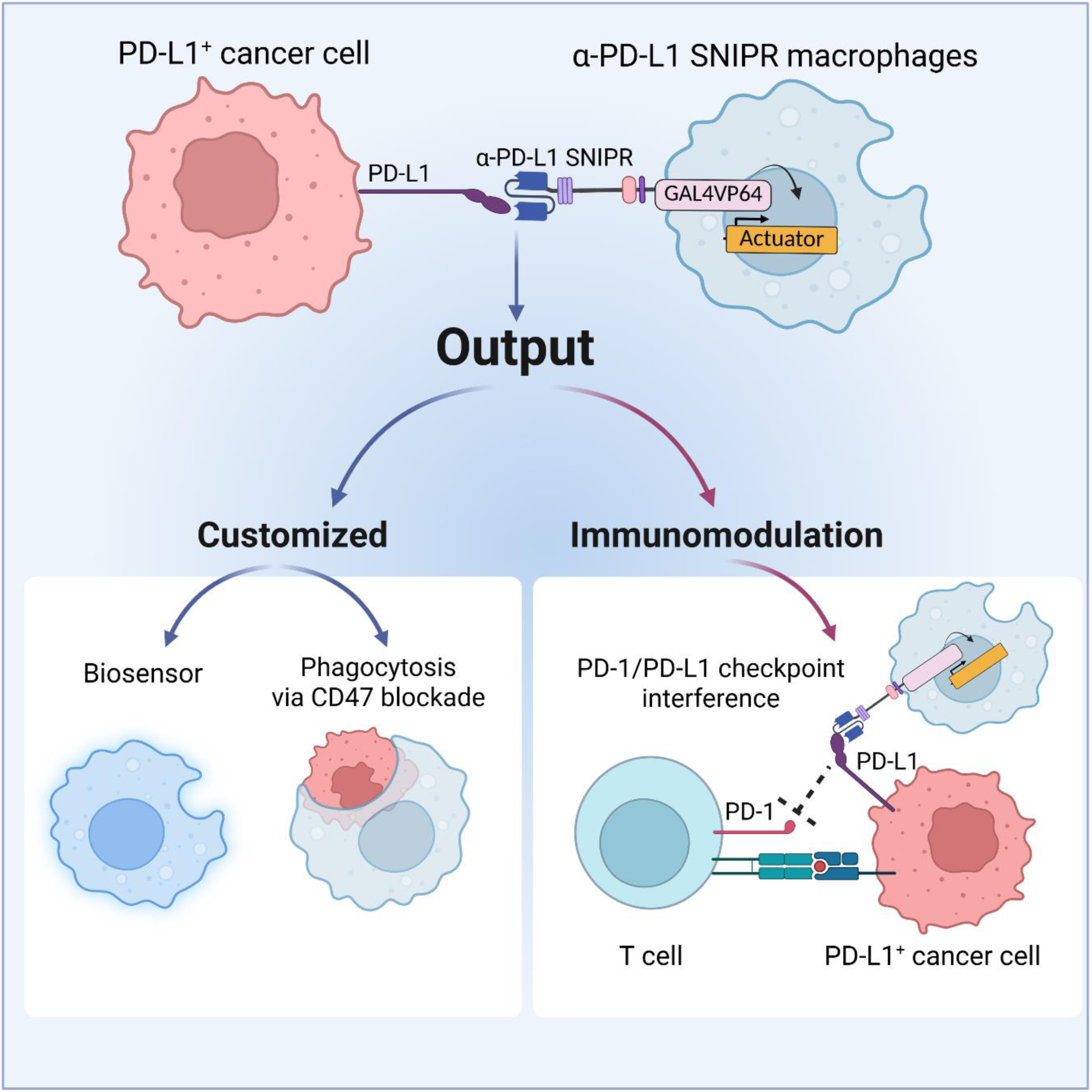

## INTRODUCTION

Synthetic biology enables the design of genetic circuits to program mammalian cell functions, or to rewire cellular response to environmental cues. In this context, such circuits typically comprise a sensor that detects a specific biomarker, such as surface ligands or secreted molecules, and an actuator that genetically encodes a defined response to that input [1,2]. This modular architecture is particularly well suited to cell-based immunotherapies, which rely on the *ex-vivo* manipulation of immune cells to augment their therapeutic function.

The intrinsic sensing-actuating capabilities of synthetic receptors made them central components of genetic circuits. Chimeric antigen receptors (CAR) and synthetic Notch receptors exemplify this paradigm by coupling defined extracellular inputs to programmable cellular responses and found successful applications in T cell-based cancer immunotherapy [3]. Despite these advances, progress in solid tumor treatment using engineered T cells has been limited by multiple challenges, including antigen escape and heterogeneity, on-target off-tumor toxicity, and impaired trafficking and infiltration [4]. These limitations have prompted the exploration of alternative strategies, including the engineering of other immune cell types.

Among these, macrophages have the advantage of naturally homing to tumor sites and represent the most abundant immune population within the tumor microenvironment (TME) across several solid tumors [5]. However, due to immunosuppressive signals in the TME and their intrinsic plasticity, tumor-associated macrophages (TAMs) are progressively skewed from a pro-inflammatory (M1-like) towards an anti-inflammatory and immunosuppressive (M2-like) state [6–9]. While pro-inflammatory macrophages are specialized in pathogens clearance and stimulation of adaptive immune responses, M2-like macrophages promote tissue repair and, in the context of solid tumors, support tumor growth, angiogenesis and metastasis formation [9,10].

Engineering macrophages with synthetic receptors has recently demonstrated therapeutic potential. CAR-expressing macrophages exhibit antigen-directed engulfment of cancer cells and an anti-HER2 CAR-M therapy has recently completed a phase I clinical trial [11,12]. Moreover, macrophages functional states can be actively reprogrammed using synthetic circuits, as exemplified by the recently described chimeric cytokine receptors (ChCR) that convert immunosuppressive cytokines such as IL-10 or TGF-β into IFN-γ signaling, thereby promoting pro-inflammatory activation [13].

Here, we present a newly designed synthetic circuit that physically links cancer cells to engineered monocytic-like cells and enhances their phagocytic capacity through target-dependent release of therapeutic molecules. To this end, we designed a synthetic Notch receptor targeting PD-L1 (α-PD-L1 SNIPR), an immune checkpoint broadly expressed by cancer cells to suppress T cell-mediated cytotoxicity.

We demonstrate that THP-1 derived macrophages engineered with α-PD-L1 SNIPR and co-cultured with SKOV-3 ovarian cancer cells undergo robust activation, triggering the expression of either a reporter gene (sensor output) or a fusion protein that blocks the CD47 “don’t eat me” signal and promotes phagocytosis *in vitro* (actuator output). Finally, we couple α-PD-L1 SNIPR signaling to endogenous macrophage transcriptional regulatory elements to restrict circuit activation in polarized macrophage.

Together, this work provides a proof-of-concept for engineering macrophages with sensor-actuator circuits capable of converting spatially constrained immunosuppressive signals into programmable therapeutic outputs.

In perspective, by interfering with an inhibitory axis expressed by both tumor and immune cells, localized release of this class of therapeutics has the potential to indirectly enhance the activity of surrounding, non-engineered immune cells within the tumor microenvironment.

## RESULTS

### Engineering and validation of a novel α-PDL1 SNIPR receptor in monocytic-like THP-1 cells

Cancer cells overexpress several membrane-bound proteins that interact with and inhibit immune cells activity. Among these, PD-L1 has gained particular attention for its ability to suppress T cell cytotoxicity through engagement of the PD-1 receptor, forming the PD-1/PD-L1 axis, the most widely targeted inhibitory checkpoint in current immunotherapeutic strategies [14–17].

Accordingly, PD-L1 represents a promising input for synthetic circuit activation for three main reasons; (i) it is highly expressed across many solid tumors [15,16], (ii) it can serve as a contact-dependent signal to rewire immune cell responses upon interaction with cancer cells and (iii) physical engagement of PD-L1 by a synthetic receptor may disrupt its interaction with PD-1 on T cells.

To connect PD-L1 sensing to a programmable cellular response, we started from the synthetic Notch (synNotch) and the more recently developed SNIPR receptor platforms, which are ideally suited due to their intrinsic contact-dependent sensor-actuator function [18–20]. Specifically, ligand binding to the extracellular domain of the receptor triggers proteolytic cleavage of the transmembrane core, resulting in release of the transcription factor that activates an ortoghonal gene expression program [18]. Moreover, the modular architecture of these receptors enables plug-and-play exchange of output domains, allowing functional adaptation to diverse downstream applications [20].

We constructed an α-PD-L1 SNIPR receptor in a lentiviral vector, by fusing an anti-PD-L1 nanobody [21], as putative extracellular domain, upstream the SNIPR core regulatory region followed by the GAL4 transcription factor as intracellular domain (**Fig. 1A**) [20]. To initially validate PD-L1 sensing, we used as read-out of the receptor activation a blue fluorescent protein (BFP) reporter under the control of a GAL4-responsive promoter (UAS-BFP). This inducible cassette was incorporated into a seperate lentiviral vector together with a second transcription unit encoding a consistutively expressed YFP reporter (**Fig. 1A**).

**Fig. 1.**
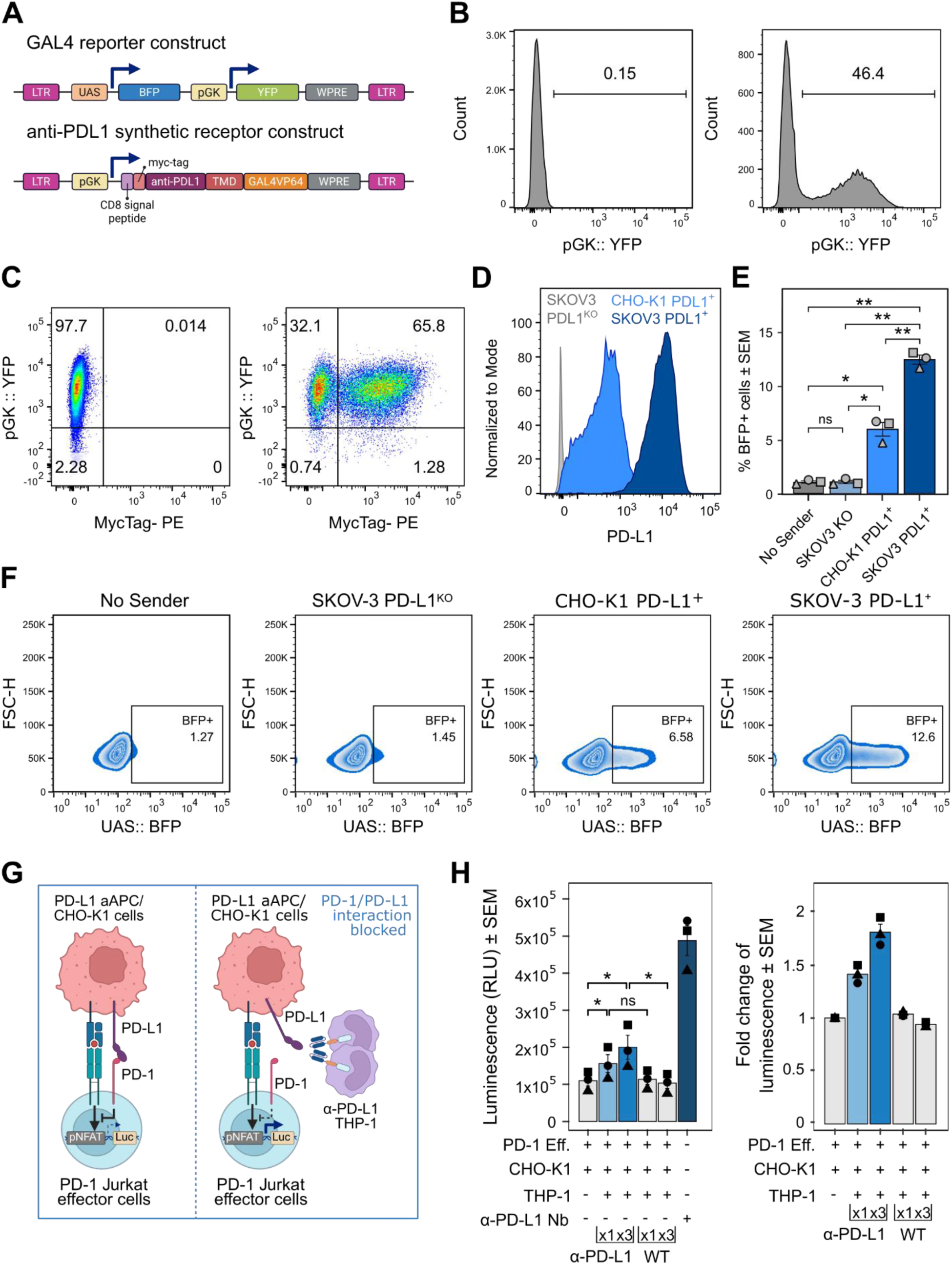
Design and validation of α-PD-L1 SNIPR in THP-1 cells. (**A**) Schematic of the lentiviral constructs encoding the α-PD-L1 SNIPR and a GAL4-responsive reporter cassette (UAS-BFP). The reporter construct includes a second transcriptional unit driving constitutive YFP expression (UAS-BFP pGK-YFP) for identification of transduced cells. (**B**) Engineering of THP-1 cells with the UAS-BFP pGK-YFP reporter construct. Flow cytometry histograms show untransduced THP-1 cells (left) and YFP expression three days after transduction, prior to cell sorting (right). (**C**) Integration of the α-PD-L1 SNIPR into sorted YFP⁺ THP-1 cells from (B). Representative flow cytometry dot plot (right) shows identification of transduced cells by YFP and Myc-tag expression three days post-transduction. (**D**) PD-L1 expression assessed by flow cytometry in PD-L1 knockout SKOV-3 cells (gray), CHO-K1 cells (light blue), and SKOV-3 cells engineered to express homogeneous PD-L1 levels (dark blue). (**E**) Fraction of α-PD-L1 THP-1 cells expressing the BFP reporter when cultured alone or after 48 hours of co-culture with the cells from (D). Statistical significance was determined using repeated-measures one-way ANOVA with Tukey’s multiple comparisons test (*adjusted p < 0.05; **adjusted p < 0.01), *n=3*. (**F**) Representative flow cytometry contour plots corresponding to the conditions shown in (E). (**G**) Schematic of the PD-1/PD-L1 signaling interference assay. PD-L1⁺ CHO-K1 antigen-presenting cells (APCs) engage PD-1 on Jurkat T cells engineered with an NFAT-responsive luciferase reporter, resulting in suppressed TCR signaling and low luciferase activity. Binding of α-PD-L1 SNIPR-expressing THP-1 cells to PD-L1 is expected to relieve PD-1–mediated inhibition and restore TCR signaling, leading to increased luciferase activity. (**H**) PD-1/PD-L1 blockade bioassay results shown as absolute luminescence (left) or fold change relative to the basal condition (co-culture of PD-1 Jurkat cells with PD-L1⁺ CHO-K1 cells). α-PD-L1 THP-1 cells were added at THP-1: CHO-K1 ratios of 1:1 or 3:1 (light blue bars). An anti-PD-L1 nanobody served as a positive control for rescue of TCR signaling (dark blue bar). Statistical significance was determined using repeated-measures one-way ANOVA followed by Fisher’s LSD test (*p* < 0.05), *n=3*.

Monocytic-like THP-1 cells were first transduced with the UAS-BFP/YFP reporter construct and YFP⁺ cells were isolated by fluorescence-activated cell sorting (FACS) (**Fig. 1A-B**). The YFP^+^ THP-1 cells were subsequently transduced with the lentiviral vector encoding the α-PD-L1 SNIPR. Inclusion of a myc-tag within the extracellular domain of the receptor enabled the identification of receptor-engineered cells. Myc-tag^+^/YFP^+^ THP-1 cells are thereafter referred to as *α-PD-L1 THP-1* cells (**Fig. 1C**, right). To evaluate circuit activation, the α-PD-L1 THP-1 cells were co-cultured with cell lines expressing different PD-L1 levels. SKOV-3 ^PD-L1+^ ovarian cancer cells and CHO-K1 ^PD-L1+^ cells exhibiting high and medium PD-L1 levels respectively, were used as test conditions whereas SKOV-3 cells in which PD-L1 was genetically ablated (SKOV-3 ^KO^) were used as a negative control [14]. For co-colture experiments, target cells were seeded one day pior to addition of the α-PD-L1 THP-1 cells and maintained in co-colture for 48 hours. We observed significant upregulation of the BFP reporter expression when α-PD-L1 THP-1 cells were co-coltured with CHO-K1 ^PD-L1+^ and SKOV-3 ^PD-L1+^ cells, whereas no activation was detected in the SKOV-3^KO^ condition (**Figure 1E-F**). Notably, circuit activation correlated with the PD-L1 expression levels on the target cells, demonstrating the sensitivity of the α-PD-L1 SNIPR receptor as a sensor of PD-L1.

### The α-PD-L1 SNIPR receptor impairs PD-L1/PD-1 interaction

Interaction between PD-L1 expressed on tumor cells and the PD-1 receptor on T cells is a key mechanism driving suppression of T cell cytotoxicity and induction of T cell exhaustion [14–16]. We reasoned that, beyond activating a synthetic circuit, engagement of PD-L1 by the α-PD-L1 SNIPR might additionally interfere with the PD-1/PDL1 axis, thereby potentially alleviating inhibitory signaling on T cells.

To test this hypotesis we employed a PD-1/PD-L1 blockade bioassay in which opposing effects of TCR activation and PD-1-mediated inhibitory signaling are quantitatively converted in the expression of a luciferase reporter (**Fig. 1G**). In this assay, disruption of PD-1/PD-L1 interaction results in enhanced TCR signaling and increased luminescence.

To evaluate the ability of the α-PD-L1 SNIPR to interfere with PD-1 binding, either unmodified wild type THP-1 cells (wt) or α-PD-L1 THP-1 cells were added to co-cultures of CHO-K1 ^PD-L1⁺^ antigen-presenting cells (APCs) and luciferase-expressing Jurkat T cells, at THP-1:APC ^PD-L1+^ ratios of 1:1 or 3:1. An anti-PDL1 nanobody was included separately as a positive control [22], to restore TCR signaling and increase luciferase activity (**Fig. 1H**). As compared to control conditions, addition of α-PD-L1 THP-1 cells resulted in a significant two-fold increase in luminescence, indicating partial restoration of TCR signaling. In contrast, addition of wild-type THP-1 cells did not significantly alter reporter activity (**Fig. 1H**).

Collectively, these results demonstrate that THP1 cells engineered with an α-PD-L1 SNIPR not only sense the presence of PD-L1^+^ cells, but also interfere with the PD-1/PDL1 axis, potentially mitigating suppressive signals of T cell activation.

### CV1-Fc improves the phagocytosis of CD47^+^ cancer cells by THP-1 derived macrophages

CD47 is a ubiquitously expressed transmembrane glycoprotein, that functions as a “marker of self”, preventing macrophage phagocytosis through interaction with signal regulatory protein α (SIRPα) expressed on myeloid cells [23]. Because CD47 is frequently upregulated on cancer cells, it provides a dominant “don’t eat me” signal that limits macrophage-mediated tumor clearance and has therefore emerged as an important target for immunotherapy (**Fig. 2A**) [24,25].

**Fig. 2.**
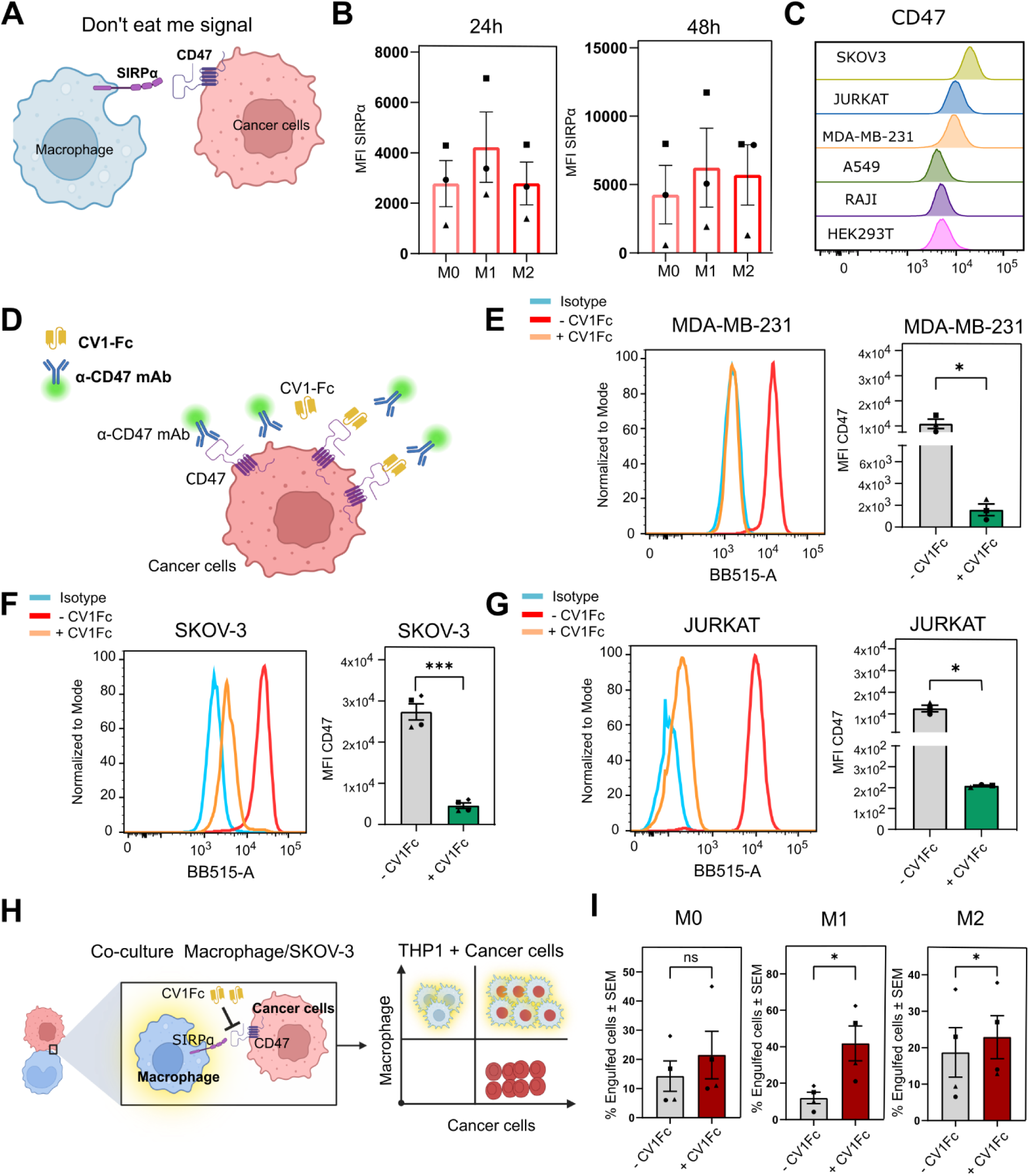
Validation of CV1-Fc–mediated enhancement of macrophage phagocytosis of CD47⁺ cancer cells. (**A**) Schematic representation of the CD47 “don’t eat me” signal. CD47 expressed on cancer cells engages signal regulatory protein α (SIRPα) on macrophages, thereby inhibiting phagocytosis. (**B)** Flow cytometry analysis of SIRPα expression on differentiated THP-1 cells following 24 and 48 hours of polarization. Bar plots indicate the median fluorescence intensity (MFI). (**C)** Flow cytometry analysis of CD47 expression across a panel of human cell lines: HEK293T (human embryonic kidney), Raji (human B-cell lymphoma), A549 (human lung carcinoma), MDA-MB-231 (human breast cancer), Jurkat (human T lymphocyte), and SKOV-3 (human ovarian adenocarcinoma). (**D)** Schematic of the CD47 competition binding assay. MDA-MB-231, SKOV-3, and Jurkat cells were incubated with CV1-Fc–enriched conditioned medium prior to staining with a fluorescently labeled anti-CD47 antibody. Binding of CV1-Fc to CD47 is expected to reduce antibody binding due to competitive, steric interference. (**E-G)** Representative flow cytometry histograms and bar plots showing CD47 staining of MDA-MB-231, SKOV-3, and Jurkat cells following treatment with CV1-Fc–conditioned medium (orange) or antibody-only control (red). Reduced antibody-associated fluorescence in the presence of CV1-Fc indicates effective competition for CD47 binding. *n=3* (**H)** Phagocytosis assay using THP-1–derived macrophages stably expressing YFP and SKOV-3 cells stably expressing mCherry. Macrophages were polarized for 48 hours and subsequently co-cultured with SKOV-3 cells for 4 hours in the presence or absence of CV1-Fc–containing conditioned medium. Double-positive YFP⁺/mCherry⁺ events represent macrophages that have engulfed cancer cells. (**I)** Quantification of cancer cell engulfment by uncommitted (M0), pro-inflammatory (M1-like), and anti-inflammatory (M2-like) macrophages. Data are presented as mean ± SEM. Statistical significance was determined using paired Student’s *t* test. *n=3*. (*p<0.05, ***p<0.001).

We reasoned that activation of the α-PD-L1 SNIPR circuit could be exploited to locally disrupt CD47-mediated inhibitory signalling in a contact-dependent manner. To this end, we selected CV1-Fc, a fusion protein composed of an affinity-enhanced SIRPα variant (CV1) fused to the Fc domain of human IgG1, which has previously been shown to mediate potent CD47 blockade, enhance macrophage phagocytosis, and activate natural killer (NK) cells, resulting in significant tumor growth inhibition in preclinical models [26,27].

Because phagocytosis is primarily executed by mature macrophages rather than circulating monocytes, we introduced a step of differentiation of the THP-1 cells into adherent macrophage-like cells with the widely used phorbol 12-myristate-13-acetate (PMA) treatment (**Fig. S1A**) [28]. PMA-treated THP-1 cells acquired macrophage morphology within 48 hours and were polarized toward either a pro-inflammatory (M1-like) or an anti-inflammatory (M2-like) state upon stimulation with lipopolysaccharide (LPS) and interferon-γ (IFN-γ), or interleukin-4 (IL-4) and interleukin-13 (IL-13), respectively (**Fig. S1A**). Polarization was confirmed by increased expression of CD80 in M1-like macrophages and CD206 in M2-like macrophages (**Fig. S1B-C**).

We then quantified SIRPα expression in uncommitted THP-1 derived macrophages (M0) as well as following 24 and 48 hours of polarization towards pro-inflammatory (M1-like) or anti-inflammatory (M2-like) states to confirm the suitability of this system for investigating CD47–SIRPα signaling. SIRPα was expressed across all phenotypes, with slightly higher levels observed in M1-like macrophages, supporting the use of this model to study CD47-SIRPα interactions *in vitro* (**Fig. 2B**). We next assessed CD47 surface expression by flow cytometry across a panel of tumor and non-tumor cell lines, including SKOV-3, Jurkat, MDA-MB-231, A549, Raji and HEK293T cells. Among these, SKOV-3 cells exhibited the highest CD47 expression levels (**Fig. 2C**).

To assess whether CV1-Fc could competitively block CD47 on tumor cells, we placed the CV1-Fc coding sequence with the IL-2 secretion peptide, under the GAL4 inducible promoter (UAS), and co-transfected HEK293T with a constitutively expressed GAL4-VP64 plasmid. 48 hours post-transfection, the supernatant containing the CV1-Fc protein was collected (**Figure S1D**) and applied to MDA-MB-231, SKOV-3, and Jurkat cancer cell lines expressing CD47. Cells were subsequently stained with a fluorescently labelled anti-CD47 antibody (**Fig. 2D**). Incubation with CV1-Fc–containing conditioned medium resulted in a significant reduction of anti-CD47–associated fluorescence across all tested cancer cell lines, indicating effective competition for CD47 binding between CV1-Fc and the anti-CD47 antibody (**Fig. 2E-G**).

We next investigated whether CD47 neutralization and Fc receptor engagement by the CV1-Fc protein enhances cancer cell phagocytosis, by co-culturing SKOV-3 cells with M0, M1-like or M2-like THP-1-derived macrophages. To quantify phagocytosis by flow cytometry, SKOV-3 cells were engineered to express a LifeAct–mCherry reporter, while THP-1 cells constitutively expressed a YFP reporter. 48 hours post-polarization, SKOV-3 cells were seeded onto polarized THP-1-derived macrophages in the presence or absence of CV1-Fc-containing conditioned medium.

Cancer cell engulfment was quantified after 4 hours of co-culture by measuring the proportion of double-positive (YFP⁺/mCherry⁺) cells (**Fig. 2H**). The presence of CV1-Fc significantly increased the fraction of engulfed SKOV-3 cells by both M1-like and M2-like macrophages (**Fig. 2I**).

Together, these results indicate that CD47-Fc fusion protein promotes cancer cell uptake independently of macrophage polarization state and highlight the potential of inducible CV1-Fc secretion as a strategy to enhance macrophage-mediated clearance of cancer cells.

### α-PD-L1 SNIPR–driven CD47 blockade enhances macrophage phagocytosis *in vitro*

Given the promising results of the CV1-Fc actuator, we incorporated this module in the α-PD-L1 circuit to enable its conditional expression upon PD-L1 detection. For this purpose, we generated a lentiviral construct containing two transcriptional units (TUs). The first TU comprises the GAL4-responsive promoter (UAS) driving the expression of CV1-Fc, followed by a P2A self-cleaving peptide and a BFP reporter. The second TU encodes a constitutively expressed LNGFR cell-surface marker to enable selection of transduced cells (**Fig. 3A**).

**Fig. 3.**
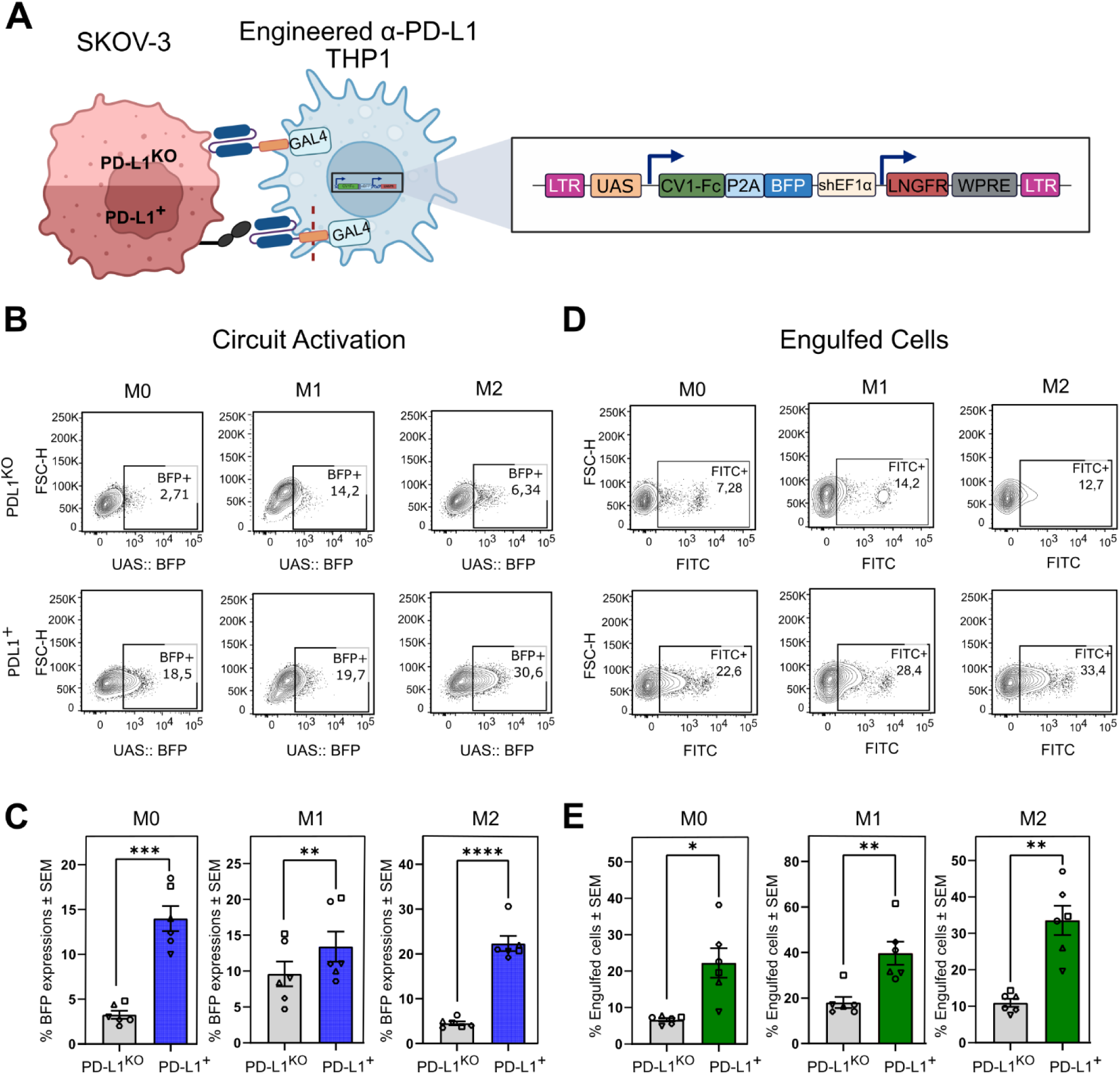
Functional validation of α-PD-L1 SNIPR–mediated induction of CV1-Fc in engineered THP-1 cells. **(A)** Schematic representation of the sensor–actuator circuit. In the presence of PD-L1–deficient SKOV-3 cells (SKOV-3 ^KO^), the α-PD-L1 SNIPR remains inactive, and CV1-Fc expression is not induced. In contrast, SKOV-3 cells expressing high levels of PD-L1 (SKOV-3 ^PDL1+^) trigger SNIPR activation and downstream CV1-Fc expression. **(B-E)** Phagocytosis assays were performed by co-culturing THP-1–derived macrophages with CFSE-FITC–labeled SKOV-3^KO^ or SKOV-3^PDL1+^ cells for 48 hours. Flow cytometry was used to simultaneously quantify circuit activation and macrophage-mediated phagocytosis. (**B-C**) Contour plots and bar plots showing BFP expression levels across uncommitted (M0), pro-inflammatory (M1-like), and anti-inflammatory (M2-like) THP-1 derived macrophages. Cells exposed to PD-L1–expressing targets exhibited increased BFP expression (blue bars) compared with ligand-negative conditions (gray bars), confirming ligand-dependent circuit activation. (**D-E**) Contour plots and bar plots showing the fraction of THP-1 derived macrophages that engulfed cancer cells identified by gating the CD11b^+^ THP-1 cells and, within this population, the FITC^+^ events Exposure to PD-L1–expressing cancer cells resulted in significantly higher phagocytic activity (green bars) compared with PD-L1–negative conditions (gray bars). Statistical significance was determined using paired Student’s *t* test. *n=6* (*p<0.05, **p<0.01, ***p<0.001, ****p<0.0001)

THP-1 cells were co-transduced with the α-PD-L1 SNIPR sensor and the responsive actuator construct. Double positive myc-tag^+^/LNGFR^+^ cells were isolated by FACS to obtain a population where both sensor and actuators were integrated (**Fig. S2A**).

Engineered THP1 cells were differentiated and uncommitted M0, M1-like and M2-like macrophages were co-cultured with PD-L1–deficient SKOV-3 cells (SKOV-3 ^KO^) or PD-L1-expressing SKOV-3 cells (SKOV-3 ^PDL1+^). Prior to co-culture, cancer cells were labelled with CFSE–FITC and seeded onto THP-1 derived macrophages. After 48 hours, cells were harvested and analyzed by flow cytometry using BFP expression as a readout of circuit activation. The macrophage sub-population was identified by gating the CD11b^+^ cells and, within these cells, the FITC⁺ events marked macrophages that had engulfed cancer cells (**Fig. S2B**). Co-culture with SKOV-3 ^PDL1+^ cells induced a significant increase of BFP expression in all differentiated macrophages. Interestingly, M0 and M2-like macrophages exhibited lower basal reporter expression in the absence of PD-L1 compared with in M1-like macrophages indicating reduced circuit leakiness (**Fig. 3B-C**). PD-L1-dependent activation of α-PD-L1 SNIPR resulted in significantly enhanced engulfment of cancer cells by engineered macrophages regardless of the polarization state (**Fig. 3D-E**). In contrast, non-engineered THP1 cells displayed comparable levels of phagocytosis toward PD-L1^+^ and PD-L1^KO^ SKOV-3 cells, indicating that enhanced engulfment was dependent on circuit activation (**Fig. S2C**).

Collectively, these results demonstrate that macrophages engineered with a sensor–actuator circuit can translate ligand-specific recognition via the α-PD-L1 receptor into a functional phagocytic output across diverse polarization states, highlighting the potential of this strategy for programmable macrophage-based interventions.

### Transcriptional control of α-PD-L1 SNIPR expression by macrophages regulatory elements

While constitutive expression of the α-PD-L1 SNIPR enables its deployment across diverse cell types, it may also lead to unintended circuit activation due to stochastic encounters with PD-L1-expressing cells. To limit unwanted induction, we leveraged the ability of synthetic, or endogenous-derived promoters to confer context-dependent control of gene expression [29–32]. We sought to restrict expression of the synthetic receptor to committed macrophages by leveraging endogenous macrophage regulatory elements (MacREs) derived from *MRC1*, *CD68*, *CSF1R* and *ITGAM* promoters [33–36]. Promoter fragments encompassing these MacREs were cloned upstream of a YFP fluorescent reporter in lentiviral vectors, followed by a constitutively expressed LNGFR marker to identify the transduced cells (**Fig. 4A**). Each MacRE construct was independently integrated into THP-1 cells, and YFP expression was quantified by flow cytometry prior to macrophages differentiation (-POL) and following polarization into uncommitted M0, pro-inflammatory (M1-like) and anti-inflammatory (M2-like) macrophages (**Fig. S3A**).

**Fig. 4.**
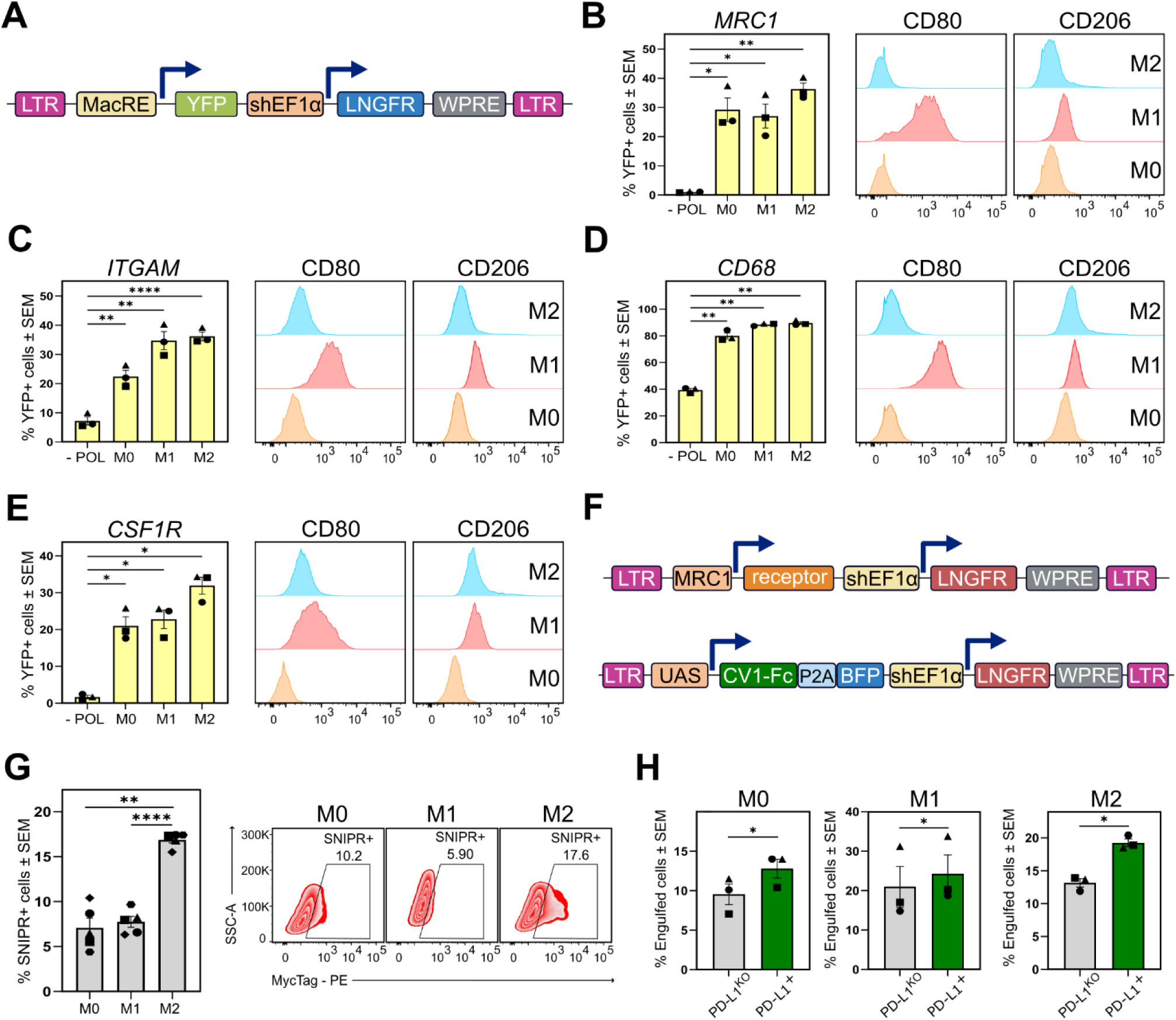
Characterization of macrophage regulatory elements (MacRE) and transcriptional control of α-PD-L1 SNIPR expression. (**A)** Schematic of the macrophage regulatory element (MacRE) reporter system. The lentiviral construct contains two transcriptional units (TUs). The first TU places a MacRE upstream of a YFP reporter. The second TU comprises a shEF1α promoter driving expression of ΔLNGFR used as a marker for transduced cells. (**B-E**) Flow cytometry analysis of MacRE activity following 48 hours of macrophage polarization. Representative histograms of the expression of phenotypic markers for M1-like (CD80) and M2-like (CD206) states are shown together with YFP expression driven by the *MRC1* (**B**), *ITGAM* (**C**), *CD68* (**D**), and *CSF1R* (**E**) promoter-derived fragments. Bar plots indicate the mean ± SEM of the fraction of YFP⁺ cells across all conditions. Statistical significance was determined using repeated-measures one-way ANOVA with Dunnett’s multiple comparisons test (*adjusted p < 0.05; **adjusted p < 0.01; ***adjusted p < 0.001; ****adjusted p < 0.001), *n=3*. **(F)** Schematic of lentiviral vectors embedding the MRC1-driven α-PD-L1 SNIPR and inducible CV1-Fc actuator constructs. Both vectors contain two TUs. The second TU is driven by the constitutive shEF1α promoter and encodes ΔLNGFR as a transduction marker. **(G)** Flow cytometry analysis of α-PD-L1 SNIPR expression driven by the *MRC1* promoter following 48 hours of macrophage polarization. Expression of α-PD-L1 SNIPR expression (left) and representative contour plots (right). Bar plots indicate mean ± SEM of the fraction of α-PD-L1 SNIPR–positive cells across polarization states. Statistical significance was determined using paired Student’s *t* test. *n=5 (*\*\*p<0.01, ****p<0.0001). **(H)** Flow cytometry analysis to quantify THP-1 that engulfed cancer cells after phagocytosis assay performed by co-culturing polarized THP-1 derived macrophages and SKOV-3^KO^ or SKOV-3^PDL1+^ cells for 48 hours. Data are presented as mean ± SEM. Statistical significance was determined using Student’s *t* test. *n=3* (*p<0.05,*p<0.01).

Across all MacREs, YFP expression was minimal at the monocytic state whereas differentiation into macrophages increased the proportion of YFP^+^ cells (**Fig. 4B-E, S3B**). The CD68-derived promoter exhibited detectable activity already in monocytes and was further activated across all polarization states (**Fig. 4D**). In contrast, *MRC1-*, *CSF1R-* and *ITGAM*-derived promoters showed little to no activity in monocytes and were induced at different levels by macrophage differentiation and polarization (**Fig. 4B-E**). Among these, the *MRC1*-derived promoter, exhibited the lowest basal activity at the monocyte-like state (-POL) with a high proportion of YFP^+^ cells following polarization, with preferential activation in the M2-like phenotype. Collectively, these results establish a framework for controlling transgene expression in macrophages using endogenous regulatory elements.

Because tumor-associated macrophages progressively acquire anti-inflammatory features, we next asked whether α-PD-L1 SNIPR expression could be preferentially restricted to M2-like macrophages to enhance circuit specificity. Based on its expression profile, we selected the MRC1 promoter and cloned it upstream of the α-PD-L1 SNIPR. The construct was integrated into THP-1 cells previously transduced with the inducible actuator module, without additional selection for cells bearing the complete circuit (**Fig. 4F**).

Engineered THP-1 cells were then differentiated and polarized, and α-PD-L1 SNIPR expression was quantified by anti-myc staining. Consistent with the YFP reporter data, receptor expression was detected upon macrophage polarization and was most pronounced in M2-like macrophages (**Fig. 4G**).

We next evaluated whether *MRC1-*driven receptor expression translated into functional circuit output. Engineered THP-1 were differentiated, polarized and co-cultured with SKOV-3 ^PD-L1+^ or SKOV-3 ^KO^ cells. While increased phagocytosis was observed across all polarization states, the effect was most pronounced in M2-like macrophages, indicating that MacRE-based transcriptional control can bias circuit activity toward defined macrophage phenotypes (**Fig 4H**).

## DISCUSSION

Macrophages possess an intrinsic tropism for tumor sites and are endowed with the ability to engulf cancer cells and cross-present antigens, thereby contributing to the initiation of adaptive immune responses. However, within the TME, immunosuppressive cytokines and dominant “don’t eat me” signals, impair macrophage phagocytosis and progressively skew their phenotype towards pro-tumorigenic states. Consequently, strategies aimed at reprogramming macrophages toward immunostimulatory functions or enhancing their capacity to directly eliminate cancer cells are actively being explored for the treatment of solid tumors.

Chimeric antigen receptor–engineered macrophages (CAR-M) have demonstrated antigen-directed engulfment of cancer cells and a favourable safety profile in early-phase clinical trial [12]. Nevertheless, these studies have thus far reported limited antitumor efficacy, potentially reflecting the predominantly cell-autonomous mechanism of action of CAR-M therapies. Given that only a small fraction of tumor-associated macrophages (TAMs) can be replaced or supplemented by engineered cells, approaches that rely exclusively on autonomous effector activity may be insufficient to broadly remodel the TME. In contrast, strategies that leverage engineered cells to locally release immunomodulatory biologics have the potential to amplify therapeutic effects by engaging the abundant endogenous macrophage population.

Consistent with this notion, blockade of the CD47/SIRPα axis has been shown to promote macrophage-mediated engulfment of cancer cells [37,38]. However, systemic administration of CD47-targeting agents is associated with dose-limiting toxicities due to the ubiquitous expression of CD47, particularly on red blood cells and platelets [39–41]. To mitigate these effects, alternative approaches have explored localised delivery of CD47-blocking molecules, including secretion by CAR-T cells [42–44]. Yet, recent studies have reported that CAR-T cells engineered to secrete CD47 blockers can themselves become targets of macrophage-mediated phagocytosis, resulting in depletion of the therapeutic cells [45,46]. In this context, macrophage-based delivery of CD47-blocking agents may offer a key advantage, as the effector molecules would be produced by cells intrinsically resistant to phagocytosis and confined to the tumor milieu. Together, these observations underscore the importance of spatially restricting the release of immunomodulatory molecules to defined cellular niches within the TME.

Synthetic Notch receptors and SNIPRs have emerged as powerful tools to achieve spatio-temporal control of therapeutic payload expression [3,20]. Leveraging their intrinsic sensing-actuating properties, we designed a SNIPR that detects PD-L1. The choice was motivated by several considerations. PD-L1 is broadly expressed across many solid tumors and is frequently associated with poor prognosis, making it a suitable marker to confine circuit activation to the TME [16,17]. In addition, physical engagement of PD-L1 by the synthetic receptor offers the opportunity to partially disrupt the PD-1/PD-L1 inhibitory axis, a major barrier to effective T-cell-mediated cytotoxicity.

Consistent with this design rationale, we demonstrated robust and target-dependent activation of the α-PD-L1 SNIPR in monocytic-like THP-1 cells, with higher induction in SKOV-3 cells exhibiting higher PD-L1 levels, mitigating an important concern related to activation outside the tumor context by cells expressing low levels of the target. Moreover, engagement of PD-L1 by the synthetic receptor partially interfered with PD-1/PD-L1 signalling, resulting in up to two-fold increase in luciferase activity in a PD-1/PD-L1 blockade assay. While a purified anti–PD-L1 nanobody produced a stronger rescue of TCR signaling, this difference is likely attributable to higher effective concentrations of the blocking agent compared with receptor-mediated blockade, which is inherently constrained by expression levels on engineered cells.

Beyond sensing and checkpoint interference, we demonstrate a proof-of-concept for coupling cancer cell recognition to enhanced macrophage phagocytic activity. To this end, we incorporated into the inducible output of the α-PD-L1 circuit, CV1-Fc, an affinity-enhanced SIRPα–Fc fusion protein previously shown to promote phagocytosis in preclinical cancer models [26,27]. We confirmed that exogenously supplied CV1-Fc enhances cancer cell engulfment by THP-1–derived macrophages in co-culture with SKOV-3 ovarian cancer cells and, importantly, we demonstrated efficient on-demand release of CV1-Fc upon α-PD-L1 SNIPR activation. Although the absolute concentration of CV1-Fc produced upon circuit activation was not directly quantified, the prolonged co-culture required for full circuit induction likely contributed to the increased phagocytosis observed, particularly in M0 and M2-like macrophages, compared to four hours of co-culture when CV1-Fc was exogenously added.

To further improve specificity and minimize unintended circuit activation, we introduced a second layer of regulation by placing α-PD-L1 SNIPR expression under the control of endogenous macrophage regulatory elements. Among the regulatory regions tested, *MRC1*-derived regulatory region showed minimal activity at the monocytic stage in THP-1 cells and preferential activation in M2-like macrophages. Consistent with this profile, *MRC1*-driven α-PD-L1 SNIPR expression resulted in enhanced engulfment of PD-L1^+^ cancer cells. The overall magnitude of phagocytosis was reduced compared with constitutive receptor expression, likely reflecting differences in promoter strength and the absence of cell sorting for complete circuit integration in the *MRC1*-driven α-PD-L1 SNIPR circuit. Nevertheless, these results illustrate the feasibility of biasing circuit activity toward specific macrophage polarization states using endogenous transcriptional control.

Overall, by combining an α-PD-L1–sensing SNIPR with a CD47-blocking actuator, we demonstrate a proof of concept in which a key cancer cell immune-evasion mechanism, PD-L1 overexpression, is exploited to override a dominant “don’t eat me” signal and enhance engulfment by THP-1–derived macrophages. By disrupting an inhibitory axis common to tumor and immune cells, localized delivery of this class of therapeutics may propagate immune activation beyond engineered cells, indirectly enhancing the activity of neighboring immune populations within the tumor microenvironment.

Future studies will be required to validate the robustness of these circuits in primary macrophages and to assess their sensitivity to additional sources of PD-L1 within inflammatory environments.

More broadly, the hierarchical regulation scheme presented here—linking sensing, decision-making, and effector delivery within engineered innate immune cells—provides a versatile platform that can be extended beyond the CD47–SIRPα axis to interrogate and manipulate other immunosuppressive pathways in the TME, serving as a flexible tool to benchmark macrophage-directed immunotherapies *in vitro*.

In summary, this study establishes a modular framework for programming macrophage behaviour through ligand-responsive synthetic circuits.

## MATERIAL AND METHODS

### Cell culture

HEK293T, MDA-MB-231, A549, THP-1, Jurkat, and Raji cell lines used in this study were purchased from the American Type Culture Collection (ATCC). The SKOV-3^KO^ and SKOV-3 ^PD-L1+^ ovarian cancer cell lines were kindly provided by the Guedan laboratory [14]. CHO-K1 ^PD-L1+^ APC and luciferase-expressing Jurkat effector cells were used as part of the PD-1/PD-L1 blockade bioassay from Promega following manufacturing istructions.

HEK293T, MDA-MB-231, A549, and SKOV-3 cells were maintained in Dulbecco’s modified Eagle medium (DMEM; Gibco) supplemented with 10% fetal bovine serum (FBS, non–USA origin; Sigma-Aldrich), 1% penicillin/streptomycin (Sigma-Aldrich), 1% L-glutamine (Sigma-Aldrich), and 1% MEM non-essential amino acids (Sigma-Aldrich). THP-1, Jurkat, and Raji cells were cultured in RPMI-1640 medium supplemented with 10% heat-inactivated FBS, 1% penicillin/streptomycin, 1% L-glutamine, and 1% MEM non-essential amino acids. All cell lines were maintained at 37 °C in a humidified incubator with 5% CO_2_.

### Cell transfection for CV1-Fc fusion protein production

HEK293T cells were transfected using PEI® transfection reagent (Polysciences) according to the manufacturer’s instructions, at a DNA (µg) to PEI (µL) ratio of 1:3. To generate conditioned medium containing the CV1-Fc fusion protein for competition and phagocytosis assays, 5×10⁶ HEK293T cells were seeded in T75 flasks in a final volume of 14 mL of complete medium. Cells were transfected with 10 µg of total plasmid DNA. Conditioned medium was collected 48 hours later, clarified by centrifugation, aliquoted, and stored at −80 °C until use.

### Lentiviral plasmid design and cloning

The sequence encoding the α-–PD-L1 SNIPR was assembled downstream of a PGK promoter. From N- to C-terminus, the construct comprises a CD8 signal peptide to ensure membrane localization; a Myc epitope tag to enable detection of surface expression of the SNIPR using a PE-conjugated anti-Myc antibody (Cell-Signaling Technology #3739) ; an anti–PD-L1 nanobody [21]; the core regulatory domain of the SNIPR receptor, consisting of the human Notch1 transmembrane domain, and juxtamembrane regions (FMYVAAAAFVLLFFVGCGVLLS-RKRRR) and a GAL4-VP64 transcriptional activator [20].

The inducible CV1-Fc actuator construct was generated starting from a plasmid kindly shared by the Ogris laboratory [27]. The coding sequence assembled downstream of UAS responsive elements included an IL-2 signal peptide, the high-affinity SIRPα fragment, CV1, fused to the Fc region of human IgG1, followed by a P2A self-cleaving peptide and a BFP reporter. To enable identification of cells harboring the complete inducible construct, a second transcriptional unit (TU) driven by a short EF1α promoter (shEF1α) and encoding a truncated low-affinity nerve growth factor receptor (ΔLNGFR) was included downstream of the actuator TU. Expression of ΔLNGFR was detected using an anti–human CD271 (NGFR) antibody (BioLegend, #345132).

Macrophage regulatory element (MacRE) reporter constructs were generated by cloning regulatory regions derived from *MRC1*, *ITGAM*, *CD68*, or *CSF1R* upstream of a YFP reporter gene [33–36]. Similar to the actuator construct, a second cassette consisting of the shEF1α promoter and ΔLNGFR was included to mark transduced cells. The MRC1:: α-PD-L1 SNIPR construct was generated by placing the α–PD-L1 SNIPR coding sequence under the control of the *MRC1*-derived promoter.

All lentiviral constructs were cloned using In-Fusion cloning (Takara Bio, #638951) into a modified pHR’SIN:CSW lentiviral backbone kindly provided by the Roybal group [20]. The GAL4 reporter construct containing UAS responsive elements driving BFP expression and a PGK promoter driving constitutive YFP expression, was also kindly provided by the Roybal group [20].

### Lentivirus production

5×10^6^ HEK293T cells were seeded from 16 to 18 hours before transfection in T75 flask in a total volume of 15mL of complete DMEM. The following day, cells were transfected with a plasmid mixture containing the pHR transfer vector, the pMD2.G envelope plasmid (Addgene #12259), and the psPAX2 packaging plasmid (Addgene #12260) at a ratio of 2:1:1. FuGENE® HD transfection reagent (Promega, #E2311) was used at a DNA (µg) to FuGENE (µL) ratio of 1:3. Within 24 hours from transfection, the culture medium was replaced to remove residual DNA and transfection reagent. Lentiviral supernatants were collected at 48 hours and 72 hours post-transfection, filtered through a 0.45-µm filter and concentrated using Lenti Concentrator (OriGene Technologies #TR30026) according to the manufacturer’s instructions. Concentrated virus was resuspended in cold DPBS, aliquoted and stored at −80 °C until use.

### Cell transduction and lentiviral titer determination

THP-1 cells were engineered using a reverse transduction protocol, in which complete medium, lentiviral supernatant, cells, and polybrene (10 µg/mL; Sigma-Aldrich) were added sequentially in this order. Transductions were typically performed in 24- or 12-well plates using 2.5–3.0 × 10⁵ cells per well in a total volume of 750 µL or 1 mL, respectively. Cells were transduced either simultaneously or sequentially, depending on the experimental design. 48 hours post-transduction, the culture medium was replaced to remove residual lentiviral particles and polybrene. Between 72 and 96 hours post-transduction, expression of circuit components was assessed by flow cytometry through quantification of YFP reporter expression for the integration of the GAL4 reporter cassette, or by staining with PE-conjugated anti-Myc antibody (Cell Signaling Technology, #3739) and Pacific Blue™-conjugated anti–CD271 (NGFR) antibody (BioLegend, #345132) to confirm the integration of sensor and actuator modules.

Lentiviral titers were determined directly in THP-1 cells seeded. Lentiviral supernatants were added to cells at dilution ratios of 1:300, 1:600, and 1:900 relative to the total culture volume. The fraction of transduced cells for each circuit module was quantified by flow cytometry as described above. Viral titers were calculated using samples yielding 20–40% transduction efficiency, according to the following formula:

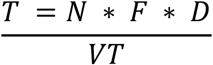

where T = Titer (TU/mL); N = Number of cells used for Titration; F = Fraction of cells successfully transduced; D = Dilution Factor; VT = Transduction Volume, mL.

### THP-1 differentiation into macrophages and polarization

THP-1 differentiation and polarization were performed using different plate formats according to experimental requirements adapting previously published protocols [28]. 4.2 × 10⁵ or 2.5 × 10⁵ THP-1 cells were seeded, respectively, onto 12- or 24-well tissue culture–treated plates in complete RPMI-1640 medium supplemented with phorbol 12-myristate-13-acetate (PMA; Sigma-Aldrich, #J63916.LB0) at a final concentration of 10 nM. Cells were incubated for 48 hours to induce differentiation before refreshing the medium to remove PMA. THP-1–derived macrophages were then kept in complete RPMI medium without any cytokine to obtain uncommitted cells (M0) or polarized toward a pro-inflammatory state (M1-like) by supplementing the medium with 10 ng/mL lipopolysaccharide (LPS; Thermo Fisher Scientific, #00-4976-93) and 20 ng/mL interferon-γ (IFN-γ; Bio-Techne, #285-IF-100) or toward an anti-inflammatory state (M2-like) by adding 20 ng/mL interleukin-4 (IL-4; PeproTech, #200-04-20UG) and 20 ng/mL interleukin-13 (IL-13; PeproTech, #200-13-10UG). Cells were incubated with polarization stimuli for 24 or 48 hours. Flow cytometric analysis was performed immediately thereafter to evaluate the expression of canonical M1 marker using Super Bright™ 702-conjugated anti-CD80 (B7-1) monoclonal antibody (BioLegend, #67-0809-42) and M2 markers with BV421-conjugated anti-CD206 conjugated to (BioLegend, #564062). SIRPα expression in THP-1 derived macrophages was measured with a PE-conjugated anti-SIRPα (CD172a) antibody (BioLegend, #372104).

### *In vitro* characterization of α-PD-L1 SNIPR activation

SKOV-3 and CHO-K1 cells were seeded one day prior to co-culture to allow adherence. Engineered THP-1 cells were subsequently added on top of the adherent cells. Different co-culture ratios were used to achieve comparable densities of adherent target cells before addition of THP-1 cells in suspension. α-PD-L1 THP-1 cells were seeded at a ratio of 2.5:1 relative to the initial number of SKOV-3 ^KO^ or SKOV-3 ^PD-L1⁺^ cells and at a ratio of 1.25:1 relative to CHO-K1 ^PD-L1⁺^ cells (Promega). After 48 hours of co-culture, cells in suspension were collected, washed with DPBS, and resuspended in FACS buffer. The fraction of cells expressing the BFP reporter within the YFP^+^ population was quantified by flow cytometry to assess circuit activation.

### PD-1/PD-L1 blockade bioassay

The capacity of the α-PD-L1 SNIPR to interfere with PD-1/PD-L1 interaction was tested by adding engineered THP-1 cells to the PD-1/PD-L1 Blockade bioassay (Promega). 3.3 × 10⁴ CHO-K1 ^PD-L1⁺^ antigen-presenting cells (APCs) were seeded 24 hours prior to co-culture. The following day, the same number of PD-1^+^ Jurkat effector cells were added to the CHO-K1 cultures together with either α-PD-L1 SNIPR–expressing THP-1 cells or wild-type THP-1 cells at THP-1: CHO-K1 ratios of 1:1 or 3:1. As a positive control for PD-1/PD-L1 pathway inhibition, an anti–PD-L1 nanobody [22] was added to the co-cultures of CHO-K1 APCs and PD-1 effector cells at a final concentration of 1 µg/mL. Luminescence was measured after 6 hours of co-culture using a GloMax® Discover system (Promega).

### CD47 Competition Assay

7.0 × 10⁴ MDA-MB-231 and 4.0 × 10⁴ SKOV-3 cells were seeded one day prior to the competition assay in 24-well plates, in a final volume of 0.5 mL of complete medium. 4.0 × 10⁵ Jurkat cells were seeded in 24-well plates on the same day of the competition assay. Cells were incubated for 1 hour at 37 °C with 0.5 mL of conditioned medium, harvested from HEK293T cells transfected to secrete CV1-Fc. Cells not incubated with CV1-Fc–containing conditioned medium were used as untreated controls. Following incubation with or without CV1-Fc, cancer cells were washed and stained with a FITC-conjugated anti-CD47 antibody (BioLegend, #11-0479-41) or a FITC-conjugated mouse IgG1 κ isotype control. Fluorescence associated with antibody binding was quantified by flow cytometry to assess competition for CD47 binding.

Staining with the same FITC-conjugated anti-CD47 antibody was also used to determine CD47 expression across different cancer cell lines such as A549, Jurkat, SKOV-3, MDA-MB-231and Raji.

### Phagocytosis assay

To assess the effect of CV1-Fc conditioned medium, 4.0 × 10⁵ THP-1 cells constitutively expressing YFP were seeded in 12-well plates in complete RPMI medium adding PMA to induce maturation to macrophages. After 48 hours, polarization stimuli were added as described above. Following an additional 48 hours, SKOV-3 cells constitutively expressing an mCherry reporter were prepared at a macrophage: cancer cell ratio of 1:3. Tumor cells were resuspended either in conditioned medium produced by HEK293T cells expressing CV1-Fc or in control RPMI medium and subsequently added to the macrophage cultures. Co-cultures were incubated at 37 °C for 4 hours, after which cells were harvested and analyzed by flow cytometry.

To assess the effect of the α-PD-L1/CV1-Fc circuit, 4.8 × 10⁵ engineered THP-1 cells were seeded in 6-well plates and differentiated with PMA as described above. After 48 hours, SKOV-3 ^KO^ and SKOV-3 ^PDL1+^ cells were prepared at a macrophage: cancer cell ratio of 1:2 and labeled with FITC-CellTrace™ CFSE Cell Proliferation Kit (Thermo Fisher Scientific, #C34554) according to the manufacturer’s instructions. Labeled tumor cells were then added to the macrophage cultures together with cytokines to induce polarization. Co-cultures were incubated at 37 °C for an additional 48 hours. Following incubation, cells were prepared for flow cytometry analysis as described below. Macrophages were stained with an PE-Cy5–conjugated anti–CD11b antibody (BioLegend, #301308), and FITC-CFSE fluorescence within the CD11b⁺ population was quantified to assess cancer cell engulfment.

### Flow cytometry and cell sorting

All cells were analyzed with a BD CELESTA™ cell analyzer (BD Biosciences). THP-1 derived macrophages were detached using sterile-filtered Accutase® solution (Sigma-Aldrich, #A6964-100ML). After 10 minutes of incubation, residual cells still attached were lifted with cell scrapers (Biosigma #010153). After detachment, cells were resuspended in 1 mL of complete medium, washed once with DPBS, and incubated with Fc Receptor Binding Inhibitor (Invitrogen, #14-9161-73) for 15 min at room temperature. Antibody staining was performed in a final volume of 50 µL of BD Horizon™ Brilliant Stain Buffer (BD Biosciences #563794) for 20 min at room temperature, washed once with DPBS, and resuspended in FACS buffer (DPBS +2mM EDTA+ 1%FBS). Population of live cells and single cells were selected according to FSC/SSC parameters. Data analysis was performed with FlowJo 10.10.0 Software (BD Life Sciences).

For cell sorting, all cells were analyzed with BD FACSMelody™ Cell Sorter. THP-1 cells were prepared in sterile conditions as described above for general flow cytometry analysis. To select the cells in which both sensor and actuators were integrated antibody staining was performed with PE-conjugated anti-Myc antibody (Cell Signaling Technology #3739) and Pacific Blue™ anti–human CD271 (NGFR) antibody (BioLegend #345132). Before sorting, cells were resuspended in 3mL of FACS medium and passed into polypropylene tubes through a 70μm cell strainer (BD Biosciences #340605). Cells were collected in complete RPMI medium, spun down, resuspended in fresh medium and seeded in multi-well plates at an approximate concentration of 5×10^5^ cells/ml.

### Statistical Analysis and figure preparation

Bar plots represent the mean of three to six independent experiments. Error bars indicate the standard error of the mean (SEM), and symbols of different shapes represent individual experimental values. Unless explicitly specified, statistical significance was determined using: two-sided paired Student’s t test for comparison between one control condition and one or two test conditions (*p < 0.05; **p < 0.01); repeated measures one-way ANOVA with Dunnett’s or Tukey’s multiple comparisons test for comparisons, respectively, among one or multiple control conditions and multiple test conditions (* adjusted p value < 0.05; ** adjusted p value < 0.01). Statistical analysis was performed using either R version 4.3.1 or GraphPad Prism 10.6.1.

For figure generation, R packages including tidyverse, rstatix, ggplot2, ggpubr, ggsignif, dplyr or GraphPad Prism were used to generate data plots and Biorender was used to generate all the illustrations. Inkscape v1.4 was used to assemble figure panels.

## Competing Interests

The authors declare no competing interests.

## Funding

The study was supported by the ERC Starting grant Synthetic T-rex (852012), the Italian Institute of Technology, and the NextGenerationEU PNRR MUR - M4C2, National Center for Gene Therapy and Drugsbased on RNA Technology” (CN00000041). L.R. was supported by the EMBO fellowship ALTF [91–2022]. M.M. was supported by the Fondazione AIRC per la Ricerca sul Cancro fellowship #31204.

## Acknowledgements

We thank Daniela Perna and Fabiana Tedeschi for technical support. Thanks to Manfred Ogris’ laboratory for providing us with the plasmid containing the CV1-Fc sequence. We thank the Smerdou group for kindly sharing the sequence of the anti-PDL1 nanobody used as putative extracellular domain of our SNIPR receptor. Thanks to the Roybal group for sending us the pHR lentiviral vectors containing the SNIPR core domain and the GAL4 responsive reporter construct. We thank Sonia Guedan for the SKOV-3 ^KO^ and SKOV-3 ^PDL1+^ cells.

## Author contributions

L.R. and V.S. conceived the project. L.R., I.DM., and M.M. designed and performed experiments and data analysis. V.S. supervised the experimental work and secured funding. L.R., I.DM., M.M and V.S. wrote the manuscript. V.S. edited the manuscript.

---

**Supplementary Fig. 1.**
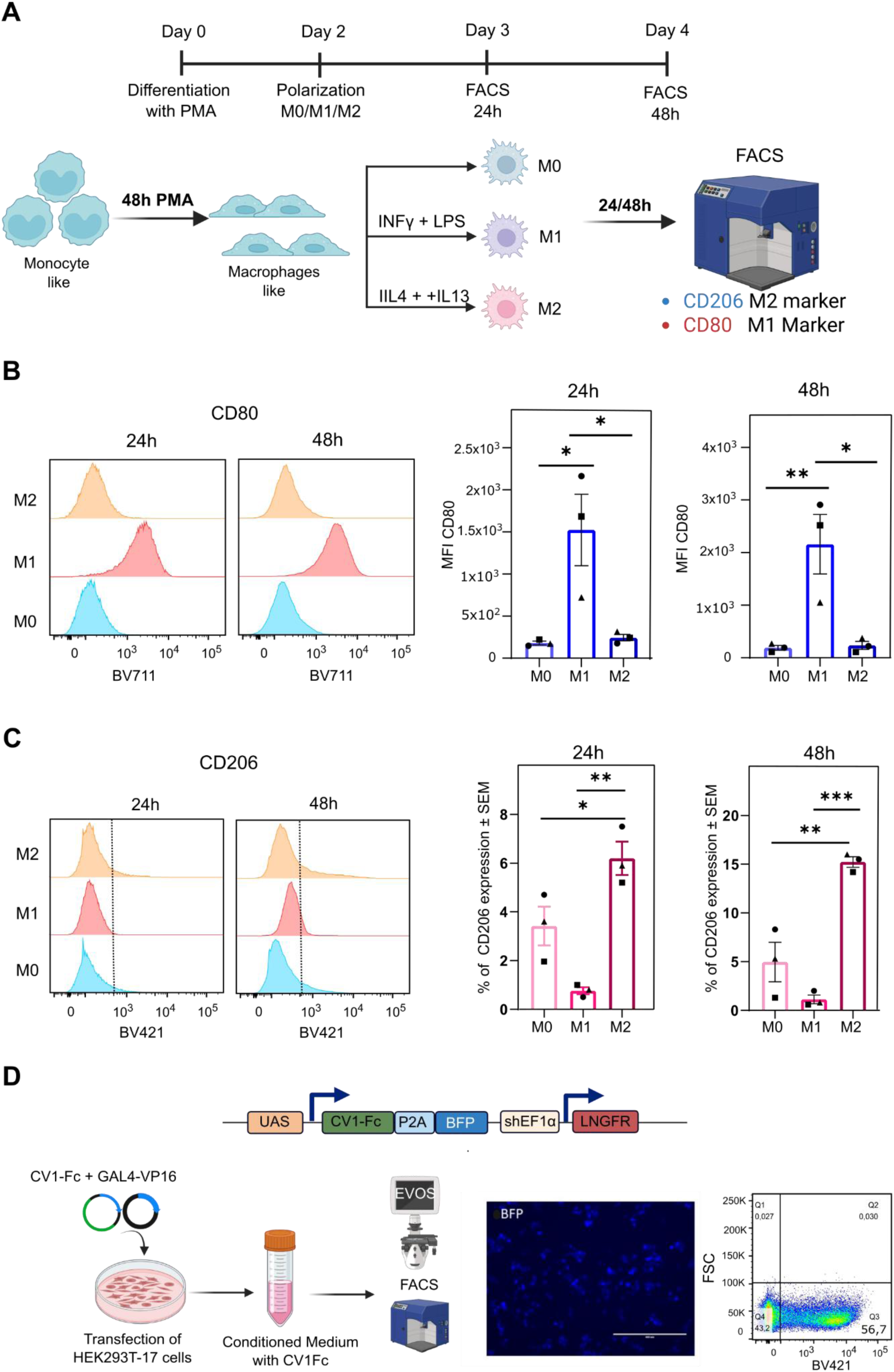
Overview of the experimental setup for THP-1 macrophage polarization and phenotypic characterization. **(A)** THP-1 monocytes were differentiated into macrophage-like cells by 48 hours of treatment with PMA and subsequently polarized toward a pro-inflammatory M1-like phenotype (IFN-γ and LPS) or an anti-inflammatory M2-like phenotype (IL-4 and IL-13). Flow cytometry analysis was performed at 24- and 48-hours post-polarization to assess surface expression of CD80 (M1 marker) and CD206 (M2 marker). **(B)** CD80 expression levels measured by flow cytometry. Bar plots indicate median fluorescence intensity (MFI) for each condition *n=3*. **(C)** CD206 expression levels measured by flow cytometry. Bar plots indicate the percentage of CD206-positive cells for each condition. *n=3*. **(D)** Production of CV1-Fc in HEK293T cells. To monitor expression of the CV1-Fc fusion protein, a BFP reporter was included downstream of the same genetic construct via a P2A self-cleaving peptide. Blue fluorescence was detected by inverted fluorescence microscopy and quantified by flow cytometry. Statistical significance was determined using repeated-measures one-way ANOVA with Dunnett’s multiple comparisons test and is presented as mean ± SEM (*p<0.05, **p<0.01, ***p<0.001).

**Supplementary Fig. 2.**
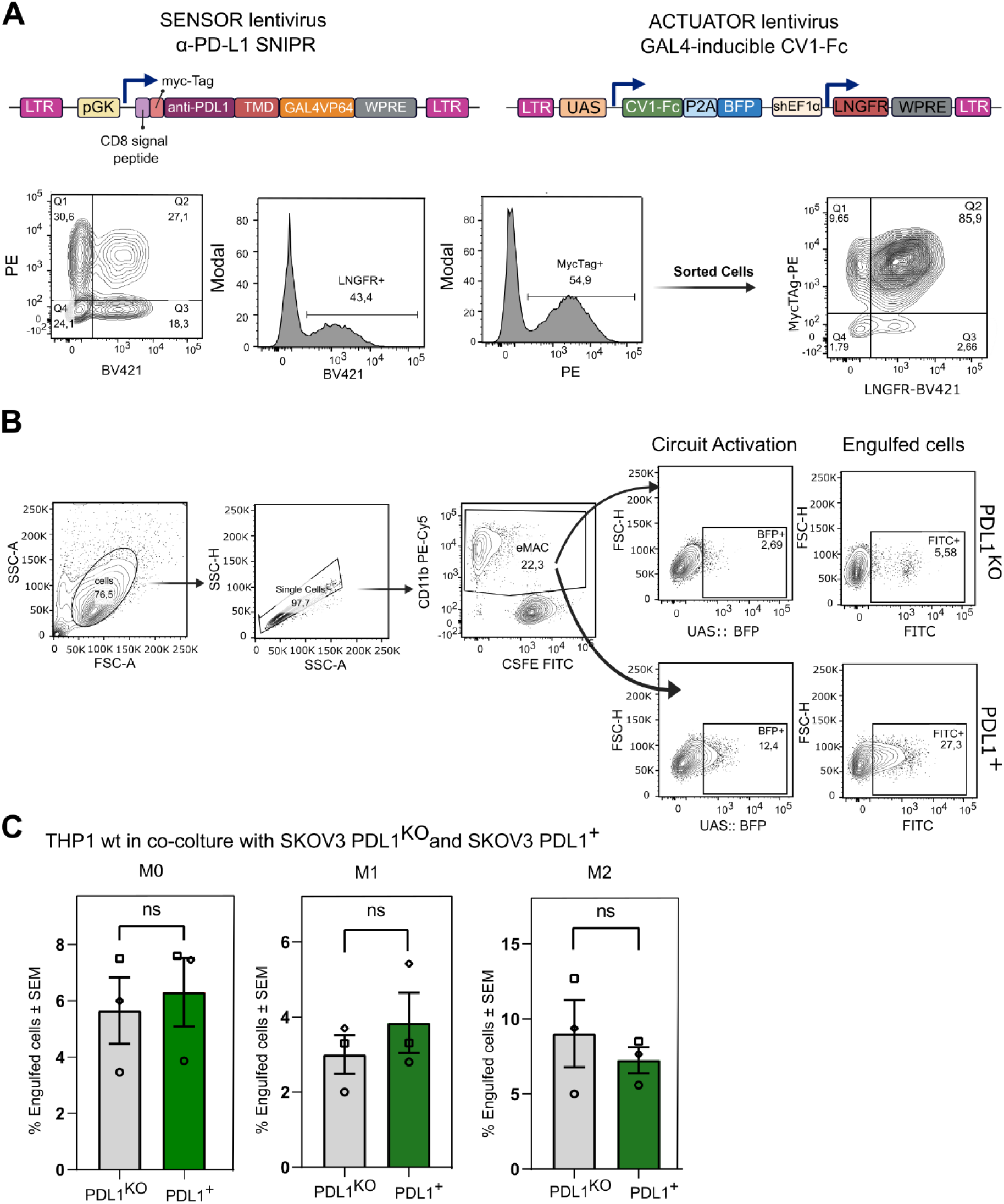
Validation of macrophage-mediate phagocytosis induced by the PD-L1 sensing. **(A)** Lentiviral constructs used to engineer THP-1 cells with the sensor–actuator synthetic circuit. The lentiviral vector (LV) was designed to generate a stable THP-1 cell line expressing both the α-PD-L1 SNIPR (sensor) and the inducible CV1-Fc actuator. The LV encoding CV1-Fc comprises two tandem transcriptional units (TUs). The first TU contains the CV1-Fc coding sequence under the control of a GAL4-responsive UAS promoter, which is activated upon ligand recognition by the synNotch receptor, resulting in CV1-Fc expression via the GAL4-VP64 transcription factor. This TU also includes a P2A self-cleaving peptide linked to a BFP reporter, enabling fluorescence-based detection of circuit activation. The second TU is driven by the constitutive shEF1α promoter and encodes truncated ΔLNGFR as a transduction marker. The α-PD-L1 SNIPR is expressed under a constitutive promoter and includes a Myc tag for detection. Following transduction, a mixed population containing approximately 27% Myc⁺/ΔLNGFR⁺ cells was obtained and subsequently sorted to yield a purified population with ∼80% double-positive cells. **(B)** Representative counterplots showing synNotch circuit activation by BFP expression and macrophage-mediated phagocytosis (engulfed cells), by GFP expression in SKOV-3^KO^ *vs* SKOV-3^PDL1+^ cells. **(C)** Phagocytosis assay performed with wild-type THP-1–derived macrophages. Following 48 hours of PMA-induced differentiation, wild-type THP-1 cells were co-cultured with SKOV-3^KO^ or SKOV-3^PDL1+^ cells for 48 hours. Bar plots show mean ± SEM of *n=3* independent experiments. No significant differences between SKOV-3^PDL1+^ cells (green bars) and SKOV-3^KO^ (gray bars). Statistical significance was determined using a paired Student’s *t* test (ns, not significant).

**Supplementary Fig. 3.**
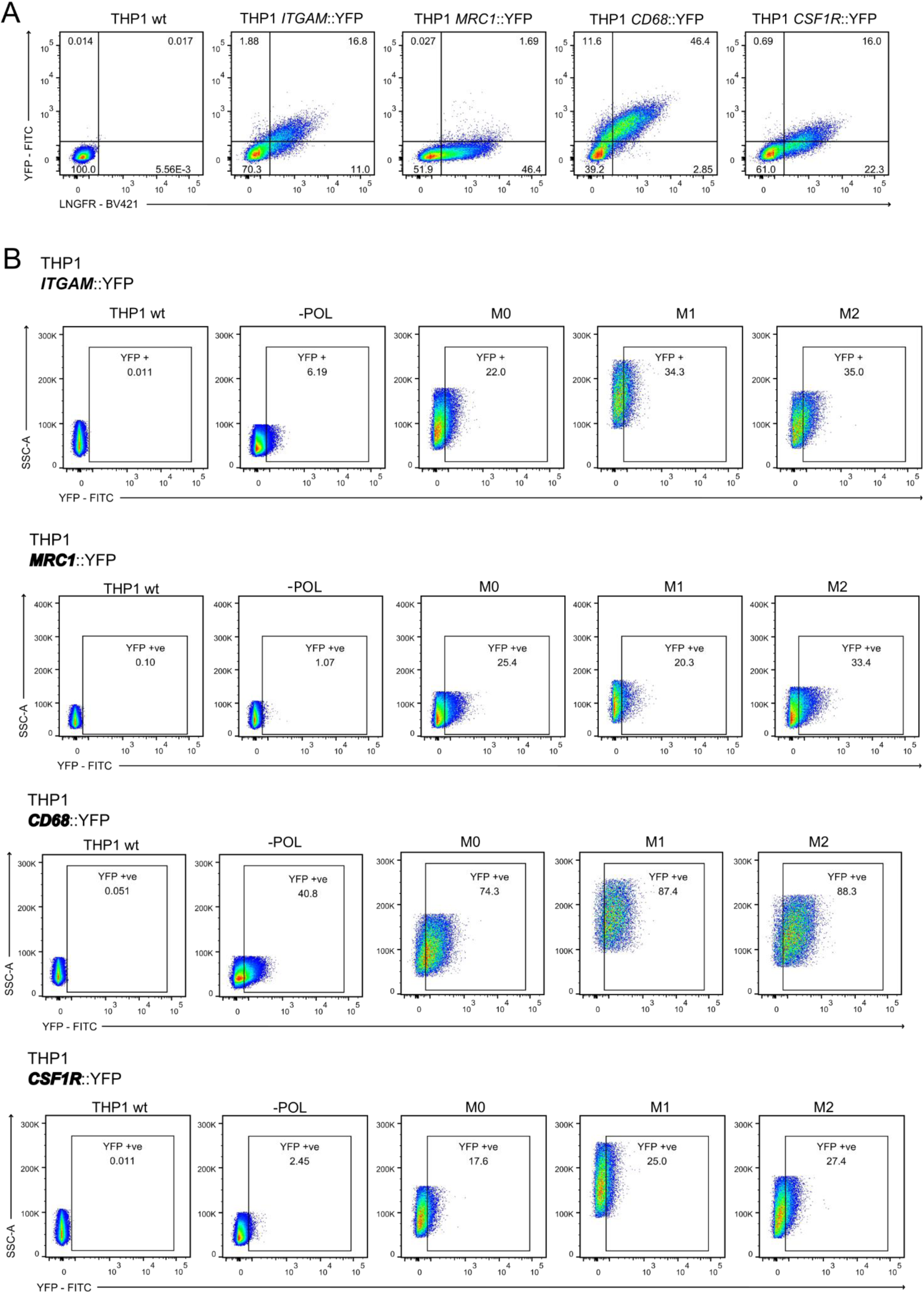
Design and validation of the MacREs reporter system. **(A)** Integration of the MacRE reporter system by lentiviral transduction was assessed by flow cytometry. Representative contour plots show the fraction of THP-1 cells expressing the constitutive ΔLNGFR transduction marker and YFP driven by the MacRE derived from *ITGAM*, *MRC1*, *CD68*, or *CSF1R* promoters, compared with untransduced control cells (left).

## Notes

### Competing Interest Statement

The authors have declared no competing interest.

